# Bacterial phylogeny explains variation in virulence but not phage efficacy *in vivo*

**DOI:** 10.64898/2026.02.05.704025

**Authors:** Sarah K. Walsh, Ryan M. Imrie, Angus Buckling, Ben Longdon

**Author notes:** Equal contribution (Angus Buckling, Ben Longdon).

## Abstract

The continuing emergence of antimicrobial resistance (AMR) poses one of the most urgent threats to public health in the 21^st^ century, causing ∼1.2 million deaths per year. A promising alternative to traditional antimicrobials is phage therapy, which has proved effective in the treatment of antimicrobial-resistant infections, including methicillin-resistant *Staphylococcus aureus* (MRSA). However, considerable variation in the outcome of phage therapy remains, with phage that show high efficacy *in vitro* often showing little to no effect when applied *in vivo.* To allow for the efficient clinical use of phage, it is vital to understand how both bacterial virulence and phage efficacy vary *in vivo* and determine how well *in vitro* and *in vivo* measures of phage efficacy correlate. Here, we infected 4,968 *Galleria mellonella* larvae with 64 phylogenetically diverse *Staphylococcaceae* isolates, both with and without co-inoculation of the bacteriophage ISP, and recorded mortality and melanisation over 24 hours. We found that bacterial virulence varied among *Staphylococcaceae* strains, and that a large proportion of this variation could be explained by the evolutionary relationships between bacteria. These results indicate that pathogen phylogeny may be a useful tool for explaining variation in both the severity of clinical infections and the virulence of novel emerging pathogens. The addition of phage significantly improved the survival of *G. mellonella* larvae, with an average 15.2% increase in endpoint survival and 10.1% reduction in endpoint melanisation across *Staphylococcaceae* strains. Phage efficacy *in vivo* showed phylogenetic repeatability, but no detectable phylogenetic heritability across bacteria species, in contrast to previous *in vitro* studies of phage infections. Concurrently, we found no evidence of a correlation between *in vivo* and *in vitro* measures of phage efficacy across bacterial isolates, demonstrating that environment plays a major role in determining the outcomes of bacteria-phage interactions. This highlights that caution should be used when extrapolating from *in vitro* measures of phage efficacy to select phages for therapeutic uses.

## Introduction

The continuing emergence of antimicrobial resistance (AMR) is one of the foremost public health threats of the 21^st^ century, a trend which is expected to continue to rise over the coming years. Recent estimates suggest ∼1.2 million deaths per year are directly attributed to AMR, increasing to ∼5 million when including deaths associated with AMR (1,2). Despite the urgent need for novel antimicrobial agents, the challenges and expense associated with antibiotic development, as well as the inevitable emergence of antimicrobial resistance, has led to few new antibiotics being approved since 2000 (3). One potential alternative to traditional antimicrobials is bacteriophage (hereafter ‘phage’), viruses which infect and kill bacteria. Phage have proven effective in the treatment of drug-resistant bacterial infections including methicillin-resistant *Staphylococcus aureus* (MRSA) (4–7) and *Pseudomonas aeruginosa* (6,7) and are being used more frequently as AMR continues to spread. To allow for the most efficient use of phage in therapeutic settings, it is important to understand phage efficacy against the target bacteria, with many studies to date investigating this using *in vitro* assays (8–11). However, while many phages demonstrate potent antibacterial activity under controlled laboratory conditions, this effectiveness is not consistently reflected in a reduction in bacterial load *in vivo* (12–16). Establishing, and ultimately improving, the extent to which *in vitro* assays can predict the therapeutic efficacy of clinically relevant phages is a key step towards increasing the efficacy of phage therapy.

Discrepancies between *in vitro* and *in vivo* phage efficacy may arise from host-mediated, bacterial, and phage-specific processes. Host-induced changes in temperature, pH, nutrient availability, and immunity can all lead to physical and chemical stresses which may alter bacterial gene expression compared to an *in vitro* environment (17,18). These changes may in turn alter bacterial susceptibility to phage infection (13,19), and explain some of the observed discrepancy between *in vitro* and *in vivo* efficacy (12–16). There are several other mechanisms that may contribute to this phenomenon, including the evolution of bacterial resistance to phage infection (20–23), differences in host immune clearance between bacteria and phage (20,24–26), and disruption from co-occurring microbes (24,27). Some procedures have been shown to improve the efficacy of phage *in vivo* prior to their application, for example, it has been demonstrated that the adaptation of phages to their bacterial hosts prior to their use in phage therapy can reduce the emergence of bacterial resistance to phage (28–30). However, whether this translates to consistent and repeatable increases in *in vivo* phage efficacy is not yet clear. Current procedures rely on *in vitro* assessments of phage efficacy prior to clinical application, yet the extent to which these assays predict therapeutic success remains unresolved.

Variation in bacterial virulence is expected to directly influence the efficacy of phage therapy *in vivo*, both by shaping host survival independently of bacterial load and by altering the immune and physiological environment in which phage-bacteria interactions occur. Thus, being able to understand how bacterial virulence can vary within and across bacterial species is key to being able to accurately assess the efficacy of phage therapy. While phage therapy has the ultimate goal of reducing or eliminating bacterial infection, there are mechanisms by which it may inadvertently increase bacterial virulence such as mediating the transfer of resistance genes (31,32). Variation in bacterial virulence across species and host environments therefore represents an additional source of heterogeneity that may contribute to inconsistent therapeutic outcomes.

Bacterial virulence is commonly mediated by specific virulence factors which may be vertically or horizontally inherited (33,34). These factors include adhesins which facilitate bacterial attachment, toxins that damage host cells and tissues, and invasins which allow bacterial invasion into host cells. For instance, *S. aureus* produces the adhesin Clumping Factor A which promotes adherence to fibrinogen on host cells (35) and the toxin Panton-Valentine Leukocidin which targets white blood cells, impairing the host’s immune response (36). Moreover, some bacteria, including *S. aureus,* deploy immune evasion factors that contribute to virulence by preventing the host’s immune response. An example of this is Protein A, which binds to the Fc region of antibodies, impeding opsonisation and phagocytosis (37). In a recent study, Laabei *et al.* explored the potential to predict bacterial virulence from genetic information by performing a genome-wide association study on 90 MRSA isolates, using phenotypic data to identify loci linked to bacterial virulence (38). This information was then used to train a machine learning algorithm which achieved over 85% accuracy in predicting the virulence of novel bacteria, suggesting that there is a wealth of genotypic data which can be used to understand and predict bacterial virulence within *S. aureus*. Such genotype-phenotype associations provide a framework for examining how variation in bacterial virulence may contribute to heterogeneity in the outcome of phage therapy *in vivo*.

*Staphylococcus aureus* is a major cause of death globally, resulting in a variety of clinical infections ranging from benign skin and soft tissue infections to severe and life-threatening systemic diseases (39,40). The recent emergence of both penicillin and methicillin resistance has led to an increase in global morbidity and mortality, with *S. aureus* infections being the second leading cause of death associated with AMR in 2019 and methicillin-resistant *S. aureus* (MRSA) alone causing more than 100,000 deaths (41). *Staphylococcaceae,* of which *S. aureus* is a member, houses several other clinically relevant bacteria (42–47), including *S. epidermidis* (42,48)*, S. saprophyticus* (49), and *S. haemolyticus* (45). The breadth of human-infective bacteria that can be found in *Staphylococcaceae* makes it an important target for phage therapy, where variation in bacterial virulence and phage efficacy across species and genera may influence clinical outcomes. It has previously been shown that the evolutionary relationships between *Staphylococcaceae* could explain a large proportion of the variation in phage susceptibility under *in vitro* conditions, indicating that many of the genetic determinants of phage susceptibility are vertically transmitted in this system (8). However, the extent to which this translates to *in vivo* phage efficacy, or how bacterial phylogeny can explain variation in bacterial virulence, remains unclear.

To this end, we used a phylogenetically diverse panel of 64 *Staphylococcaceae* isolates (spanning 2 genera and 17 species) and the bacteriophage ISP (Intravenous Staphylococcal Phage) to investigate bacterial virulence and phage efficacy *in vivo* using *Galleria mellonella* larvae. ISP is a double-stranded DNA virus closely related to *Staphylococcus* phage G1 (50,51) which has previously been used for the treatment of clinical infections (12,52). *G. mellonella* have gained traction as a model organism for bacterial infections due to their affordability, ease of use, and complex innate immune response which shares homology with vertebrate systems, containing both cellular and humoral responses (53–56). In *G. mellonella*, a central component of the innate immune response is melanisation, which contributes to pathogen encapsulation, generation of reactive oxygen species, and wound healing (57–60). Activation of this pathway results in the accumulation of melanin within the larvae, providing a visible indicator of immune activation and host damage over time. *G. mellonella* have previously been used to study various aspects of *staphylococcal* infections including infection dynamics (61–63) and biofilm formation (64–66), as well as being used as a model organism for the treatment of bacterial infections with both antibiotics (67–70) and phage (71–74).

Here, we show that bacterial virulence varies widely across *Staphylococcaceae* and that a large proportion of this variation is explained by the evolutionary relatedness of bacterial strains. *In vivo* phage efficacy also varies substantially across bacterial strains. However, whilst patterns of phage efficacy are repeatable across related bacteria, there was no evidence for the bacterial phylogeny explaining variation in phage efficacy in vivo. Consistent with this, we find no detectable correlation between *in vivo* and *in vitro* measures of phage efficacy across species. Together, these results indicate that the bacterial phylogeny may act as a useful tool for the prediction of phage efficacy and infection virulence, and that the host environment can decouple *in vivo* and *in vitro* measures of efficacy, indicating that caution should be used when selecting phages for therapeutic use based on evidence from *in vitro* assays.

## Methods

### *Staphylococcaceae* isolates

The *Staphylococcaceae* panel comprises 64 strains of *Staphylococcaceae*, representing a broad phylogenetic and geographic range of hosts. These strains span 2 genera and 17 species that were estimated to have last shared a common ancestor ∼122 Ma (75). Multi-locus sequencing typing (MLST) revealed that 5 Clonal Complexes (CC) of *S. aureus* were represented in this panel, including major complexes CC1 and CC8 (Table S1).

To minimise differences arising from laboratory adaptation, isogenic populations of each *Staphylococcaceae* strain had previously been isolated and stored in 25% glycerol at -80°C. When required, isolates were cultured by inoculating 10mL of high-salt LB Miller broth (Formedium) in a sterile 30mL glass universal with a scraping of the frozen culture and incubating at 37°C, 180rpm overnight. To ensure that any observed differences in virulence or susceptibility to ISP were not a function of bacterial density, calibration curves using optical density (OD) and colony forming units (CFU)/mL were generated and used to dilute or concentrate each strain to a final concentration of 1×10^10^ CFU/mL prior to use.

### Bacteriophage

An isolate of ISP was kindly provided by Jean-Paul Pirnay and Maya Merabishvili at the Queen Astrid Military Hospital (Brussels, Belgium). ISP was propagated on the *S. aureus* strain EOE35 (hereafter referred to as the ‘propagation host’) and extracted using chloroform. The number of infectious phage present was quantified using plaque assays performed as previously described (8).

### Galleria mellonella

Final instar *Galleria mellonella* larvae were obtained from LiveFoods Direct (Sheffield, UK) and stored at 4°C in the dark to prevent pupation. Prior to injection, larvae were sorted into groups of 13 and stored in petri dishes at 4°C until required (maximum of 24 hours). Larvae with signs of injury, melanisation, or that were noticeably smaller or larger than average were excluded. The order in which individual larvae were assigned to each treatment was randomised to account for any subconscious biases in selection.

Body size has been shown to influence infection outcomes in *G. mellonella* (76). To provide a measure of body size while minimising handling stress, a pilot set of 120 larvae were weighed and imaged (as described below) to establish the relationship between larval mass and pixel area. A linear model revealed a strong positive relationship between pixel area and weight (Pearson’s r = 0.94, p <0.001; Figure S1). Experimental larvae were then assigned weights based on the average of their pixel areas across images taken at each experimental timepoint, and weight was included as a covariate in all subsequent analyses.

### Inoculations

Bacteria were diluted to 1×10^10^ CFU/mL and phage to 1×10^13^ PFU/mL. For the without phage treatment, the appropriate bacterial strain was diluted 1:2 with LB Miller Broth to ensure an equivalent total injection volume and bacterial dose. 10uL of the diluted bacteria was injected into the last left proleg of each larva using a 50µL Hamilton syringe (Hamilton, Nevada, USA) for a final dose of 1×10^8^ bacteria. For the with phage treatment, bacteria were diluted 1:2 with phage immediately prior to injection, for a final dose of 1×10^8^ bacteria and 1×10^11^ phage (MOI of 1000). These doses were chosen based on pilot data which showed sufficient continuous variation in bacterial virulence and phage efficacy across strains at these doses within an experimental timeframe that minimised censoring due to larval pupation.

Thirteen larvae were injected for each treatment to account for excessive post-injection bleeding, which occurred in a subset of individuals, and could alter the received dose of bacteria or phage. Larvae were left to recover for 15-minutes on filter paper before the twelve showing the least bleeding were transferred to each well of a 12-well plate and incubated at 37°C in the dark. Mortality – defined here as larvae being sessile and non-responsive to stimuli – was measured once an hour from 0- to 12-hours post infection (hpi), and at a final additional timepoint of 24 hpi. To minimise block effects and reduce experimental noise, bacterial treatments were randomised across the 10-day injection schedule. For each bacterial strain, three replicate groups of larvae were prepared, and on each day, twenty randomised combinations of bacterial strain and replicate were selected for injection. Both with and without phage treatments were performed on the same day for a given replicate. All inoculations were performed by a single individual to avoid introducing inter-operator variability.

Melanisation was quantified by photographing each plate at each hourly time point using a Panasonic Lumix GH3 camera fitted with a G Vario 14-45 mm f/3.5-5.6 lens. The camera was fixed with a tripod at 50cm vertically from the plate surface, and RAW images were taken at a fixed focal length of 35 mm. Plates were evenly lit by overhead diffusion lighting, and imaged on an 18% neutral grey background, which provided a consistent reference for white balance and exposure correction during image analysis. The bottom of each plate well was covered with a matte green cardboard disc created using a 2.2 cm diameter hole punch (Hobbycraft, UK) to allow for the separation of each larva from the background image by chroma keying.

Following image acquisition, RAW (.RW2) images were processed using a custom Python script. RAW files were first demosaiced and converted to RGB, then white balance and exposure were normalised using the neutral grey card included in each image. Plate wells were identified by an automated circle-finding algorithm: green-dominant pixels were first identified by comparing RGB channel intensities, generating a binary mask corresponding to the green well inserts. This mask was refined using iterative morphological erosion and dilation to remove noise and fill gaps. For each well, a fixed-radius circular mask was placed on an approximate starting centre coordinate, and the location of this mask was varied iteratively in all cardinal and diagonal directions – first in large (20px), then progressively smaller steps (10px, 5px, 2px, 1px) – to maximise overlap between the circular mask and the green-dominant pixels of the well. Circular masks were then applied to extract individual cropped images of each well.

Within each cropped well image, larvae were segmented from the green well inserts by identifying red-dominant (i.e., low-green) pixels. The largest single connected component in each well was assumed to be the larva and retained, and the boundaries of the larva mask were refined using morphological filtering before being applied. False colour validation images (Figure S2) were generated for each plate showing the positions of well and larva masks. These images were manually inspected to detect any poor segmentation with the automated script. In a small number of cases, haemolymph leakage onto the green well insert caused larva masks to include a large proportion of dark-green background. In these cases, the larva masks were manually adjusted in image editing software (Affinity Photo 2) to remove the well background. For each segmented larva, the total number of pixels was recorded as a proxy for larval size, and mean RGB values were extracted and converted to a greyscale luminance value, with melanisation then calculated as 1 − *luminance*. The minimum and maximum observed melanisation values were then used as bounds to define 0-100% melanisation within the context of this experiment.

To ensure any experimental observations were not due to pre-existing illness, injection trauma, or inoculum contamination, no-injection and injection controls (LB Miller broth or ISP only) were included on each day of injections. Control plates throughout the experiment showed no detectable mortality and little evidence of melanisation (Figure S3).

### *In vitro* measures of phage efficacy

To compare phage efficacy measured *in vivo* and *in vitro*, we used previously collected optical density (OD) and qPCR data (8). Briefly, for OD assays, each *Staphylococcaceae* isolate was incubated for 24 hours at 37°C in the presence or absence of bacteriophage ISP. Phage-treated cultures had final concentrations of 5x10⁴ CFU/mL bacteria and 2.5x10³ PFU/mL phage (multiplicity of infection, MOI = 0.05), while control cultures contained bacteria only. Three biological replicates were performed for each treatment condition. Following incubation, optical density was measured at 600 nm using a MultiSkan Sky Microplate Spectrophotometer (ThermoFisher). *In vitro* phage efficacy was quantified as the proportional reduction in bacterial growth over 24 hours due to phage exposure, calculated as:

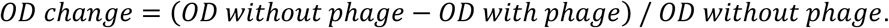

This metric was used as a measure of bacterial growth suppression by phage under controlled *in vitro* conditions.

For qPCR, infections were set up identically to the OD assays described above. Following the 24-hour incubation, each sample was heat treated at 90°C for 10-minutes to inactivate the bacteria before viral DNA was extracted using Chelex 100 (Merck Life Sciences) and mechanical disruption. qPCR was performed on each of the samples using an Applied Biosystems StepOnePlus system with a Sensifast Hi-Rox Sybr kit (Bioline). Cycle conditions were as follows: initial denaturation at 95°C for 120 seconds, then 40 cycles of 95°C for 5 seconds, and 60°C for 30 seconds. ISP was measured using the following primers: forward, 5’- CCTGTACCGGCTTGACTCTC -3’; reverse, 5’- AGCTACAACCGAGCAGTTAGA -3’. For each sample, two technical replicates of the qPCR reaction were performed and mean viral C_t_ values from technical replicate pairs (C_t:24_) were normalised to an initial dose of ISP (C_t:0_) and converted to log_10_ fold change in viral load using the 2^-ΔCt^ method, where ΔCt = C_t:24_ – C_t:0_.

### Inferring the host phylogeny

The methods used to infer the host phylogeny have been described in detail elsewhere (8). Briefly, whole genome sequences were collected for eight previously sequenced strains from NCBI (Table S1), and the remaining 56 sequences were obtained through whole genome sequencing. Library preparations and sequencing were performed by MicrobesNG (Birmingham, UK) and a core genome of the isolates was constructed by identifying orthologous genes shared between all 64 *Staphylococcaceae* isolates. The final core genome alignment contained 102 genes.

Ultrametric phylogenetic trees were constructed using BEAST v1.10 (77). Sequence alignments were fitted to HKY substitution models using relaxed uncorrelated molecular clock models, gamma distributions of rate variation, and constant population size coalescent priors (77). Separate substitution models and molecular clocks were fitted to 1^st^/2^nd^ and 3^rd^ codon positions, to reflect differences in selective constraint (78). Two independent MCMC chains were run until convergence was achieved, with a burn-in of <10%. Convergence of all parameters was checked using Tracer v1.6 (79).

### Statistical analysis

To allow for the joint analysis of mortality and melanisation – which show different non-linear relationships with time – non-linear least squares models were used to provide summary parameter estimates which could be used as response variables in phylogenetic mixed models. These models were fitted separately to data from each plate of 12 larvae, yielding n = 3 independent replicates (plates) for each unique combination of bacterial isolate and ISP. Mortality data followed typical survival curve dynamics, but with curves frequently seen to reach a plateau below 100% mortality within the observational window (Figure S4), indicating the presence of surviving individuals (i.e., a “cure fraction”). In some cases, mortality was low or absent during the 0-12 hour window but present at 24 hours, giving rise to two equally plausible survival curves: one in which the observed 24-hour mortality represents the upper asymptote of mortality (referred to here as the “cured” assumption), and one in which it marks the onset of a larger, unobserved increase in mortality beyond the observational window (referred to here as the “unobserved” assumption). All results and analyses presented below use parameter estimates inferred under the cured assumption, and parallel analyses performed using the unobserved assumption yielded qualitatively similar results. Melanisation tended to increase rapidly in the first few hours of infection, then slow to a plateau by the end of the observation window (Figure S5). Michaelis-Menten curves were chosen to summarise these dynamics, as they provide two parameters analogous to those of survival curves: in both cases, the fitted models provided estimates of an upper asymptote (maximum mortality or melanisation) and a T_50_ describing the time to half-maximal response. For both melanisation and mortality, the fitted models provided a close description of the observed data (Figure S6).

Phylogenetic generalised linear mixed models were then used to investigate the extent to which the evolutionary relationships between bacteria could explain variation in the maximum and T_50_ parameter values for mortality and melanisation, and the change in these values due to the inclusion of ISP. The structures of these models were as follows:

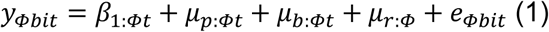

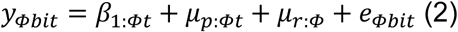

In these models, *y_Φbit_* is the parameter estimate of trait *t* (maximum mortality or melanisation; T_50_ mortality or melanisation; or larval weight) in infection condition *Φ* (with phage; without phage; effect of phage Δ) for the *ith* biological replicate of bacterial strain *b*. The fixed effect *β*_1_ is the intercept for each trait; the random effect *μ_p_* represents the effect of host phylogeny under a Brownian model of evolution; *μ_b_* represents a categorical bacterial-strain-specific effect that is independent of phylogeny and explicitly estimates the non-phylogenetic component of trait variance. Lastly, *μ_r_* represents a random effect of experimental block, and *e* is the model residuals. All random effects and residuals were assumed to be Gaussian distributed with a centred mean of 0.

The proportion of between-strain variance that can be explained by the bacterial phylogeny was calculated from model (1) using the equation 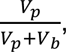 which is equivalent to phylogenetic heritability or Pagel’s lambda (80,81). Here, *V_p_* and *V_b_* represent the phylogenetic and bacteria strain-specific components of trait variance respectively (82). The repeatability of traits within-bacterial-strains after accounting for any phylogenetic variance was calculated as 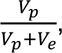 where *V* and *V* are the phylogenetic and residual variance components from model (2). Inter-strain correlation coefficients were calculated from model *V_p_* matrices 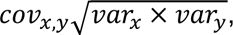 and correlation slopes as 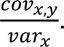 For parameters on an unbounded scale (correlation coefficients, slopes), effects were considered significant when the 95% credible intervals (calculated here as 95% HPD intervals) did not include zero. For proportions of variance (heritability, repeatability), effects were considered non-zero when the lower bound of the 95% credible interval exceeded a value of 0.05. This threshold was chosen to avoid treating effects whose posterior distributions allocate a high probability mass near zero as meaningful.

Models were fitted using the R package MCMCglmm (83) in R v4.5.2 (84), and were run for 13 million MCMC generations, sampled every 5,000 iterations with a burn-in of 3 million generations. Parameter expanded priors were placed on the covariance matrices, resulting in multivariate F distributions with marginal variance distributions scaled by 1,000. Inverse-gamma priors were placed on the residual variances, with a shape and scale equal to 0.02. To ensure the model outputs were robust to changes in prior distribution, models were also fitted with flat and inverse-Wishart priors, which gave qualitatively similar results. Parameter estimates reported are means of the posterior density, and 95% credible intervals (CIs) were taken to be the 95% highest posterior density intervals.

All scripts and data used in this analysis are available at https://github.com/Sarah-K-Walsh/2026-In-Vivo-Phage-Efficacy.

## Results

### Virulence in *G. mellonella* is a heritable trait across *Staphylococcaceae*

To investigate whether the evolutionary relationships between bacteria could explain variation in their *in vivo* virulence and susceptibility to phage, we experimentally infected 4,608 *G. mellonella* larvae with 64 *Staphylococcaceae* isolates in the presence or absence of bacteriophage ISP, and recorded mortality and melanisation over the following 24 hours (129,024 total observations).

*G. mellonella* larvae showed considerable variation in their mortality and melanisation rates across the bacterial strains tested (Figures 1 and 2, respectively). Some bacterial strains caused mortality in all hosts within 8 hpi (e.g., *S. aureus* B142S1 and DEU2) while for other strains, a subset of hosts consistently survived infection beyond the observational window (e.g., *S. aureus* ASARM74 and EOE003). Similarly, some strains induced high levels of melanisation with either a rapid (e.g., *M. sciuri*) or gradual onset (e.g., *S. shleiferi spp coagulans*) and others caused little melanisation (e.g., *S. aureus* EOE041 or *S. aureus* SaTPS3072).

**Figure 1:**
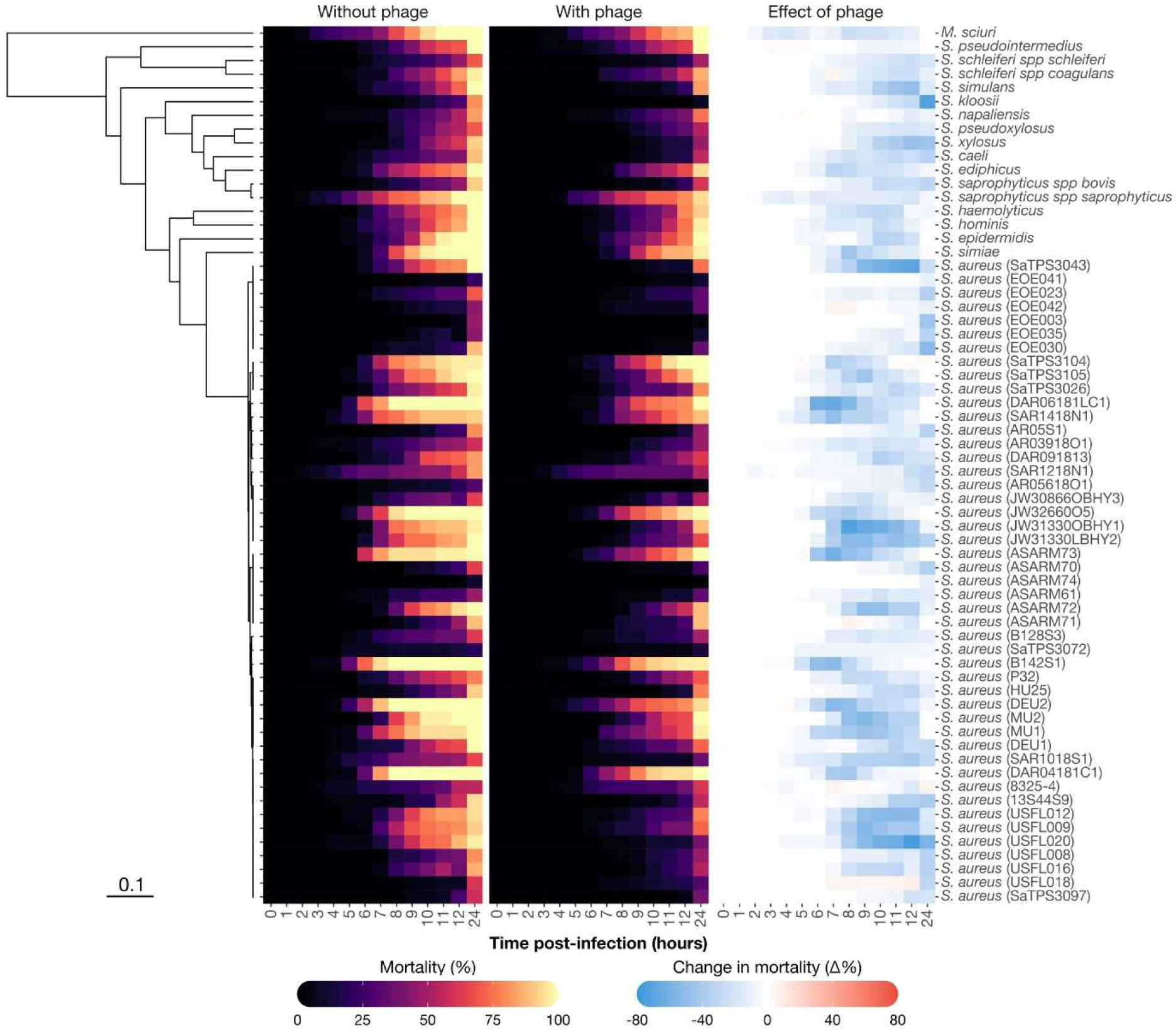
Mortality of *G. mellonella* larvae following *Staphylococcaceae* infection with and without co-inoculation of ISP. Heatmaps show percentage mortality over time (‘Without phage’ and ‘With phage’ panels) and change in percentage mortality with ISP (‘Effect of phage’ panel) for 64 *Staphylococcaceae* strains infecting *G. mellonella* larvae. Values shown are means across three biological replicates (plates of twelve individuals). A maximum clade credibility bacterial phylogeny is shown on the left, with a scale bar indicating substitutions-per-site.

**Figure 2:**
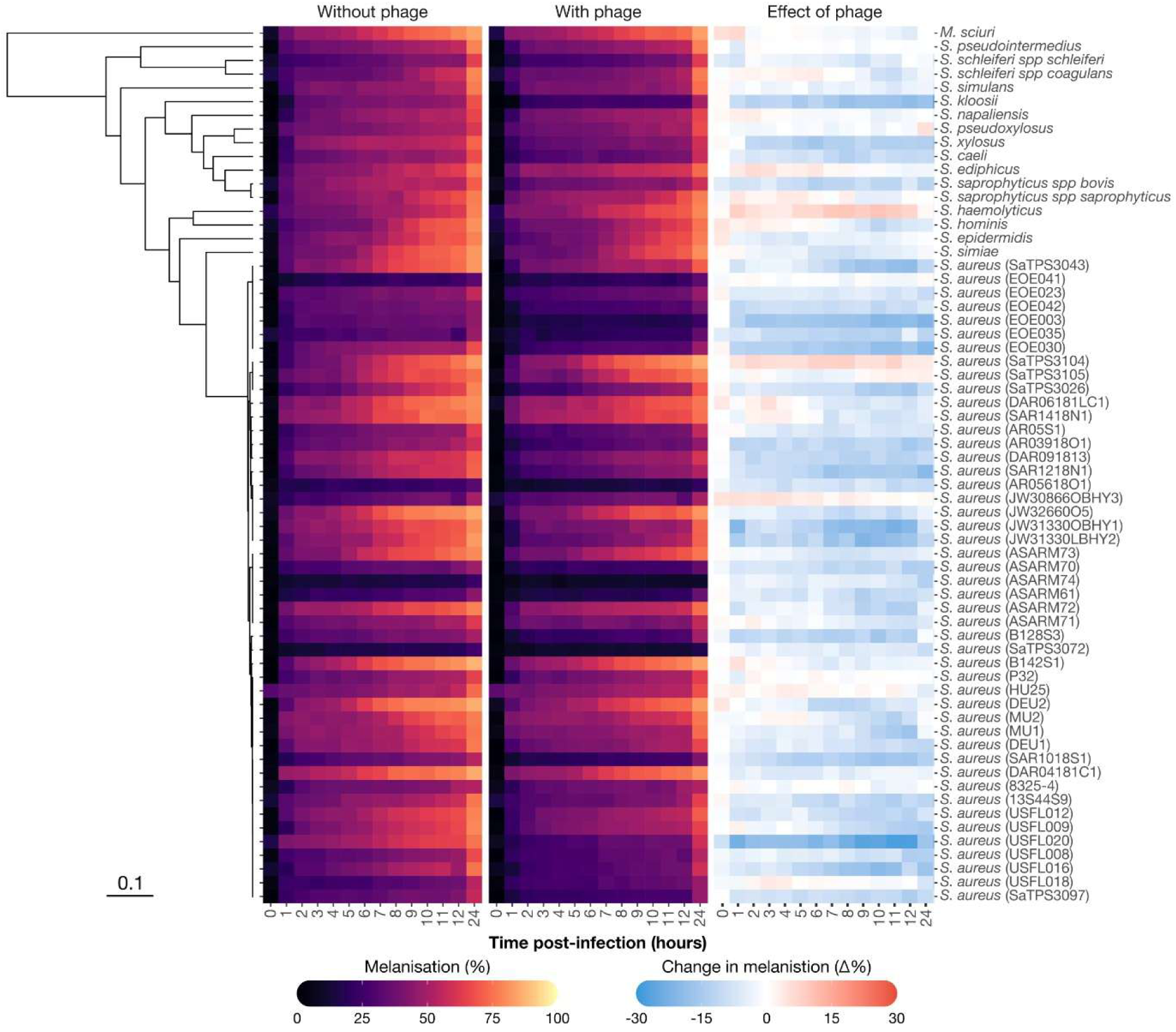
Melanisation in *G. mellonella* larvae following *Staphylococcaceae* infection with and without co-inoculation of ISP. Heatmaps show percentage melanisation over time (‘Without phage’ and ‘With phage’ panels) and change in percentage melanisation with ISP (‘Effect of phage’ panel) for 64 *Staphylococcaceae* strains infecting *G. mellonella* larvae. Values shown are means across three biological replicates (plates of twelve individuals). A maximum clade credibility bacterial phylogeny is shown on the left, with a scale bar indicating substitutions-per-site.

Non-linear least squares models were used to summarise the dynamics of both mortality and melanisation over time, describing them in terms of their maximum (upper asymptote) and T_50_ (time to ½ of maximum). Across bacterial strains, mean maximum mortality was 92.6% (95% CIs: 54-100%) with a mean T_50_ of 10.2 hours (95% CIs: 9.3-10.9 hours), suggesting that *Staphylococcaceae* infections are, in general, highly virulent in *G. mellonella*. Concurrently, the mean across-strain maximum melanisation was high (67.7%, 95% CIs: 25-100%) with a mean T_50_ of 2.1 hours (95% CIs: 0.2-4 hours) indicating a rapid onset of the melanisation response. When these parameters were further analysed in phylogenetic mixed models, all four showed evidence of phylogenetic structure across *Staphylococcaceae* (Figure 3, ‘Without phage’), with credible estimates of phylogenetic heritability detected for maximum mortality (0.63, 95% CIs: 0.06-1.00), maximum melanisation (0.88, 95% CIs: 0.24-1.00), and melanisation T_50_ (0.51, 95% CIs: 0.38-0.63). A credible estimate of repeatability (0.86, 95% CIs: 0.66-0.99), but not phylogenetic heritability, was detected for mortality T_50_. Together, this suggests that a large proportion of the variation in virulence across *Staphylococcaceae* in *G. mellonella* larvae can be explained by the bacterial phylogeny.

**Figure 3:**
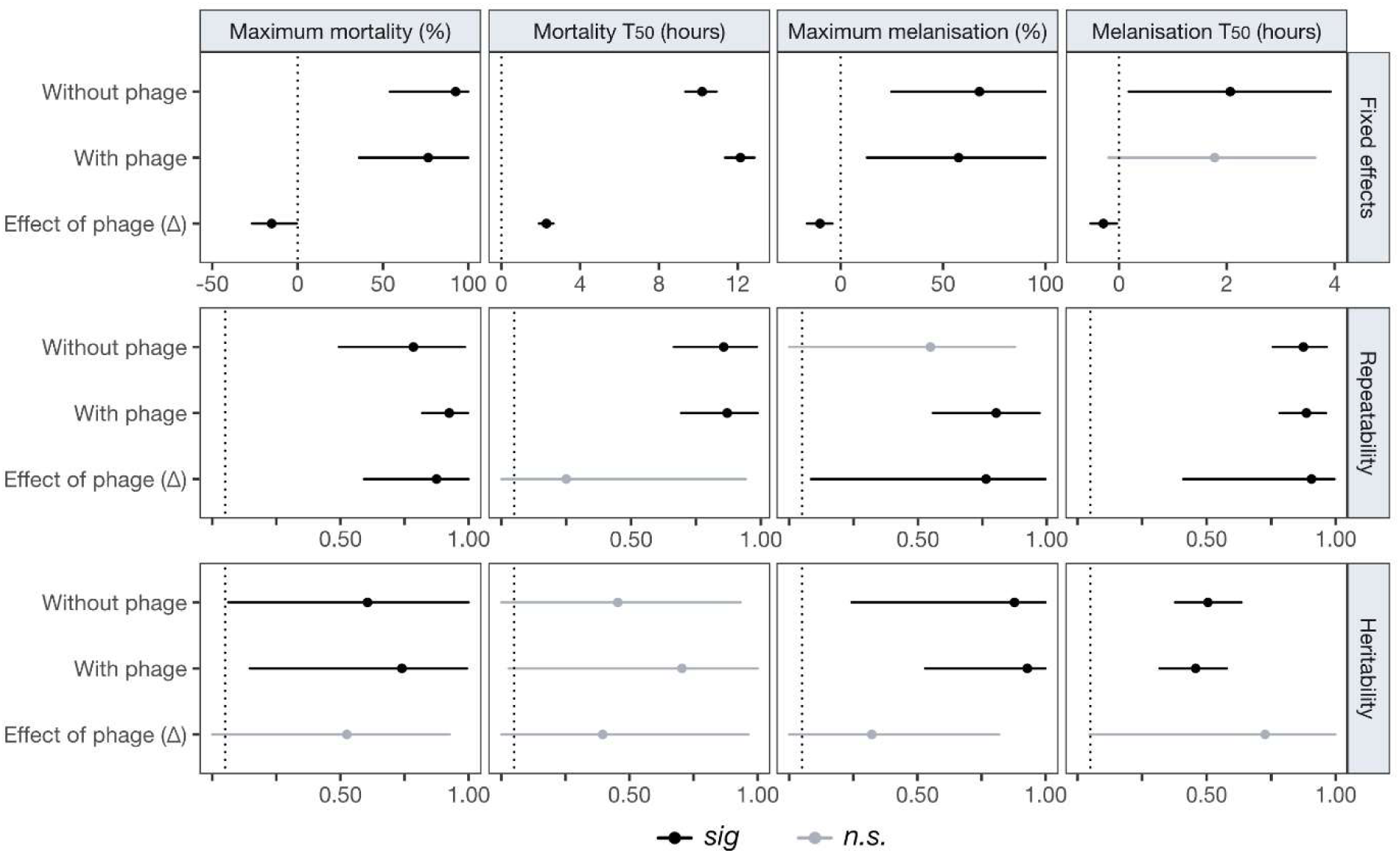
Estimates of fixed effects, repeatability, and phylogenetic heritability for mortality and melanisation parameters across *Staphylococcaceae* strains. Panels show the posterior means (points) and 95% highest-posterior density (HPD) intervals (lines) for phylogenetic mixed model estimates of inter-strain means (fixed effects), repeatability, and phylogenetic heritability of four virulence parameters: maximum mortality (%), mortality T_50_ (hours), maximum melanisation (%), and melanisation T_50_ (hours). Heritability estimates are taken from models of structure (1), while fixed effect and repeatability estimates are taken from models of structure (2). Dotted lines are used to highlight 0 (fixed effects) and 0.05 (repeatability, phylogenetic heritability). Credible estimates, whose HPD intervals do not contain these values, are highlighted in black.

### Phage efficacy varies *in vivo* and is partly explained by bacterial phylogeny

Co-inoculation of bacteria and ISP generally led to a decrease in both mortality (Figure 1) and melanisation (Figure 2), although considerable between-strain variation was again evident, with some strains showing large changes in infection dynamics with phage (*e.g*., *S. aureus* USFL020 and *S. kloosii*) and others appearing largely unaffected (*e.g*., *S. aureus* 8325-4 and *S. napaliensis*). When comparing the across-strain means with and without phage treatment, the credible intervals of most mortality and melanisation parameters overlapped (Figure 3, except for mortality T_50_). However, when analysed as a change in parameter value with the addition of phage (in effect treating the data as paired) credible across-strain mean effects of phage treatment were detected for all four parameters. These estimates suggest that, on average, the inclusion of ISP caused a 15.2% decrease in maximum mortality (95% CIs: 1-27%); an increase of 3.2 hours to mortality T_50_ (95% CIs: 2.5-3.8 hours); a decrease in maximum melanisation of 10.1% (95% CIs: 4-16.4%); and a decrease of 0.29 hours to melanisation T_50_ (95% CIs: 0.03-0.53 hours).

The phylogenetic signal in bacterial virulence remained consistent with the addition of phage (Figure 3, ‘With phage’), with little change in the rank-order of virulence among bacterial strains in the presence or absence of ISP (Figure S7). In contrast, no credible estimates of phylogenetic heritability were detected for the effect of ISP on any mortality or melanisation parameters (Figure 3, ‘Effect of phage’), although the phage-induced change in three of these parameters did show credible estimates of repeatability: maximum mortality (0.88, 95% CIs: 0.61-0.99); maximum melanisation (0.76, 95% CIs: 0.08-1.00); and melanisation T_50_ (0.91, 95% CIs: 0.44-1.00). Together, these results provide limited evidence that the efficacy of ISP tends to be more similar among closely related *Staphylococcaceae* strains.

### *Staphylococcaceae* virulence traits are correlated across-strains

To investigate whether mortality and melanisation parameters were correlated across *Staphylococcaceae* strains, inter-strain correlations were estimated for each pair of parameters using phylogenetic mixed models (Figure 4). Positive correlations were detected between maximum melanisation and melanisation T_50_ (R = 0.48, 95% CIs: 0.16-0.81), and between maximum melanisation and maximum mortality (R = 0.51, 95% CIs: 0.16-0.81), indicating that infections which induce greater endpoint melanisation tend to also induce longer melanisation responses and have higher endpoint mortality rates. Negative correlations were also detected between mortality T_50_ and maximum melanisation (R = - 0.40, 95% CIs: -0.04, -0.75), and between mortality T_50_ and maximum mortality (R = -0.51, 95% CIs: -0.19, -0.80), suggesting that infections which cause rapid mortality are often also those with the highest endpoint mortality rates and melanisation responses. No credible correlations were detected between maximum mortality and melanisation T_50_, or between mortality T_50_ and melanisation T_50_. The strength of all detected correlations remained consistent in the presence or absence of ISP.

**Figure 4:**
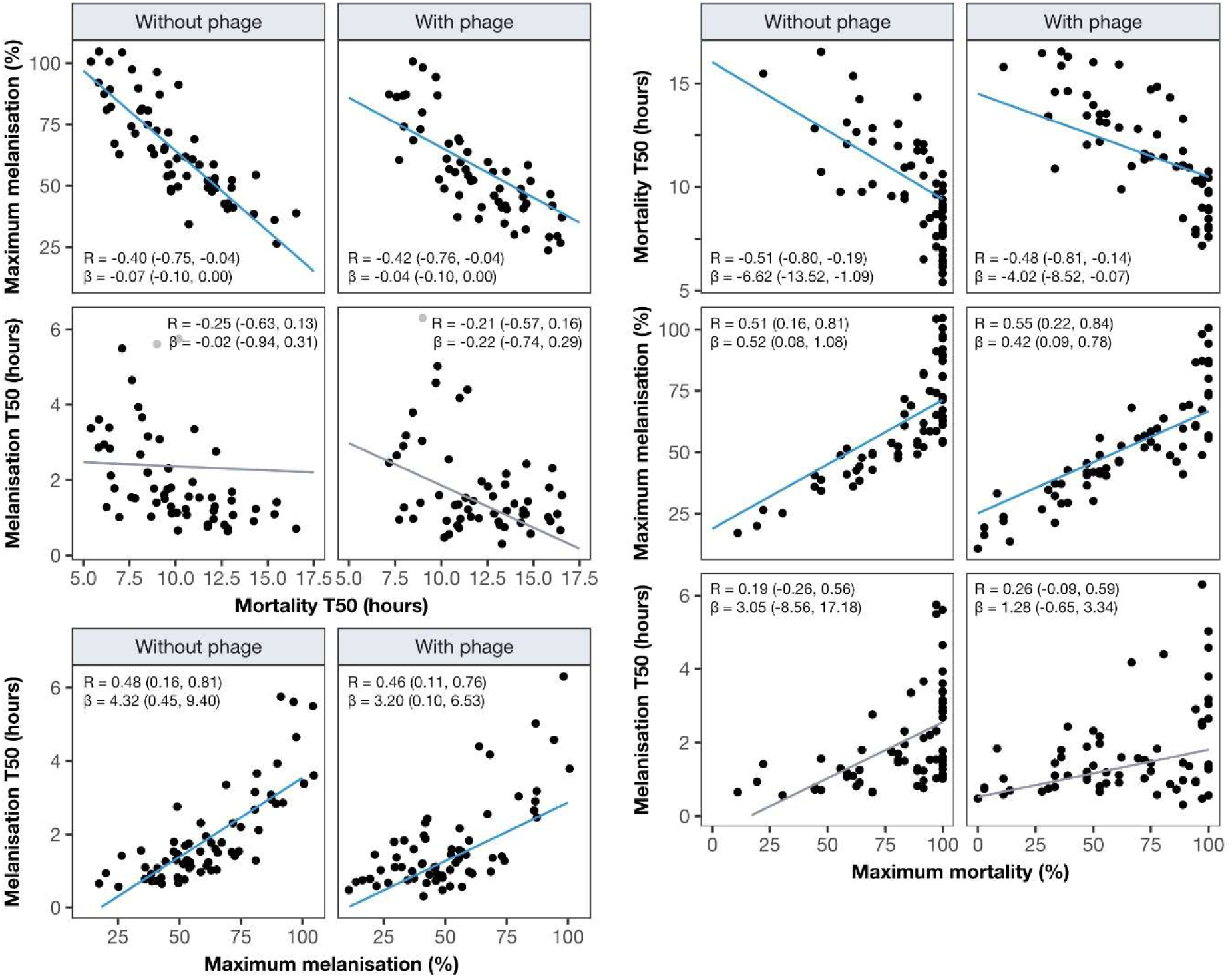
Inter-strain correlations between mortality and melanisation parameters in the presence and absence of ISP. Plots show the inter-strain correlations of each pair of mortality and melanisation parameters, with points indicating the mean value of each *Staphylococcaceae* strain. Trendlines, correlation coefficients and slopes are taken from phylogenetic mixed models of structure (1), with trendlines whose coefficient and slope estimates do not include zero highlighted in blue.

### No detectable correlation in ISP efficacy *in vivo* and *in vitro*

Finally, we investigated whether ISP efficacy *in vivo* – measured as changes in mortality and melanisation parameters with phage treatment – showed any detectable correlation with efficacy *in vitro*, using previously collected data on ISP virulence (optical density assays) and replication (qPCR) (8). No credible estimates for correlation coefficients were found for any pairs of *in vivo* and *in vitro* measures of efficacy (Figure 5), indicating that the host environment greatly alters the relative susceptibility of *Staphylococcaceae* strains to phage treatment.

**Figure 5:**
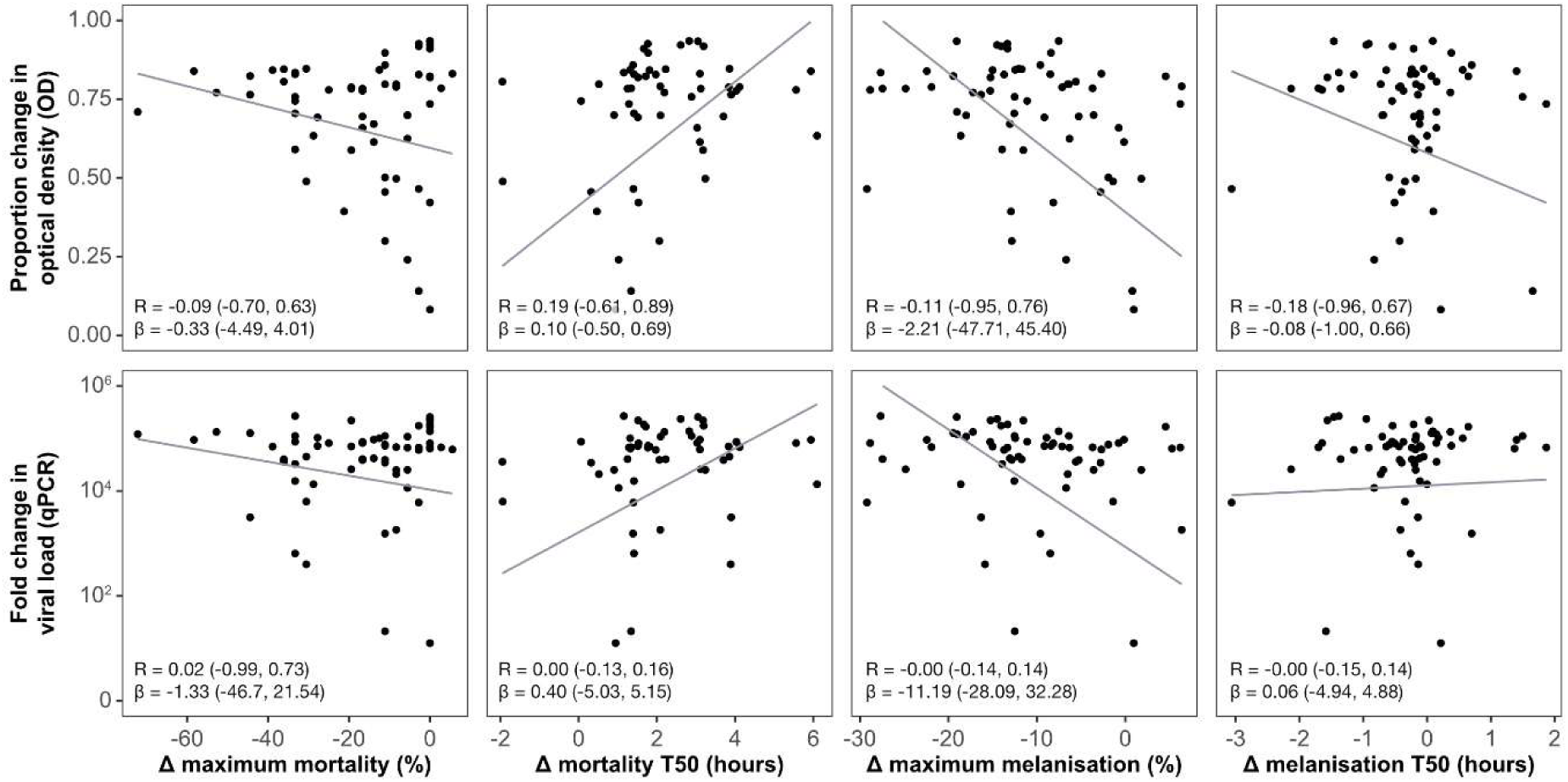
Inter-strain correlations between *in vivo* and *in vitro* measures of ISP efficacy. Plots show the inter-strain correlations of each pair of *in vivo* (maximum mortality, mortality T_50_, maximum melanisation, and melanisation T_50_) and *in vitro* (OD, qPCR) measures of ISP efficacy, with points indicating the mean value of each *Staphylococcaceae* strain. Trendlines, correlation coefficients and slopes are taken from phylogenetic mixed models of structure (1).

## Discussion

Here, we have examined the extent to which the evolutionary relationships between bacteria can explain variation in virulence and phage susceptibility *in vivo*. We found that bacterial virulence varied across strains, and that the bacterial phylogeny could explain a large proportion of this variation. Co-inoculation of ISP significantly reduced the maximum mortality, speed of mortality, and maximum melanisation experienced during infection, and variation in these components of phage efficacy showed evidence of repeatability across bacterial strains (but not detectable phylogenetic heritability). Finally, we found no correlation between the effect of phage *in vitro* and *in vivo,* suggesting that the outcome of phage infection *in vivo* is largely influenced by the host environment.

Previously, we showed that the evolutionary relationships between bacterial hosts can explain variation in susceptibility to phage infection *in vitro* (8). However, here we have shown that *in vivo,* the relationship between bacterial relatedness and susceptibility to phage is less pronounced, with only credible estimates of repeatability for phage efficacy across *Staphylococcaceae.* This suggests that the additional sources of variation imposed by the *G. mellonella* host environment – such as immunity, local pH, or nutrient availability – may alter the role of bacterial phylogeny in explaining phage infectivity *in vivo*. Correspondingly, no detectable inter-strain correlations were detected between any *in vivo* and *in vitro* metrics of phage efficacy, indicating that the host environment can act as a strong disruptor of *in vitro* phylogenetic signal. This pattern is consistent with a “layered” model of infection outcomes, where multiple layers of variation operating across biological scales combine to produce a measurable effect, with higher-order layers having the potential to mask the signal in lower-order layers (85). The first step in tailor-made phage therapy is often the *in vitro* assessment of a clinical isolate against a panel of phages to determine which one will be the most effective, a process which assumes that the efficacy of a phage *in vitro* will directly translate into *in vivo* efficacy. This reliance on *in vitro* assays to select phage for therapy may explain some of the observed inconsistency in *in vivo* phage efficacy; with phage showing highly variable success rates in treating similar infections (12).

Host immune responses can influence bacterial populations and, by extension, phage dynamics, through various mechanisms including direct effects on bacterial or phage survival, and indirect effects on bacterial susceptibility to phage infection (86,87). The innate immune system has been shown to play a role in the clearance of bacteriophage *in vivo* (88–90), including neutralisation by the complement system (91–93) or phagocytosis (89,94). In our data, we see several instances in which the addition of phage led to a marked early increase in *G. mellonella* survival compared to the no-phage treatment, with no corresponding improvement in endpoint survival (*e.g.*, *S. aureus* DAR06181LC1), a pattern that may be consistent with phage clearance within 24 hours of inoculation. Repeated or sustained phage dosing may modify the dynamics observed here by maintaining phage densities above thresholds required for productive infection and *in situ* amplification. This would be consistent with pharmacokinetic and pharmacodynamic studies which show threshold phage densities must be sustained at the infection site to enable *in situ* amplification and bacterial clearance (95,96). However, it has also been observed that repeated dosing of phage can lead to lower serum phage titres and increased host immune clearance (97,98). In this system, repeated dosing was not explored because it would require multiple injections per larva, which can introduce injection-related mortality and confound survival outcomes in *G. mellonella*. Further characterisation of the mechanisms by which the host immune system interacts with phage, and how this impacts phage persistence and replication within hosts, will be important for understanding the complex dynamics of phage treatment *in vivo*.

Most studies to date have examined how host relatedness can explain variation in infection traits such as host susceptibility (99–105), with few exploring the role of pathogen phylogeny in explaining pathogen traits (106,107). While differences in bacterial virulence have been associated with different phylogenetic clades (108–113), including within *S. aureus* (114–119), formal investigations into the influence of phylogeny on bacterial virulence remain rare. Here, we find that the evolutionary relationships between bacteria can explain a considerable proportion of their virulence *in vivo*, with closely related strains of bacteria tending to show similar levels of virulence. This similarity in virulence between closely related strains is likely due to similarities in the genetic factors underpinning infection, including similarities in the ability of the bacteria to persist and replicate within a host (120–122); bacterial virulence factors (123,124); and the expression of virulence factors, for example, differences in quorum sensing (125,126). Conversely, we do observe some instances where closely related species show large differences in their virulence (*e.g*., the *S. aureus* strains EOE041, EOE023, and EOE003), suggesting that there are instances where Brownian evolutionary processes alone cannot explain variation in virulence. This may reflect evolutionarily recent changes in the genetic components which influence virulence (*e.g.,* single nucleotide polymorphisms in virulence factors (127–129)), or the loss or gain of genetic components in a non-vertically inherited way (*e.g.*, horizontal gene transfer leading to the acquisition of a novel virulence factor (130–133)). Improving our understanding of how the relatedness of bacterial strains can explain their virulence, and the taxonomic scales at which this relationship breaks down, may improve our ability to predict bacterial virulence, disease outcome, and manage treatment plans in a clinical setting.

Together, our results demonstrate that bacterial phylogeny is an important explainer of pathogen virulence *in vivo,* with implications for our ability to predict both the virulence of clinical infections and of novel emerging pathogens. Furthermore, we highlight that *in vivo* phage efficacy shows no detectable correlation with *in vitro* measures of phage efficacy, suggesting alternative approaches may be required to assess phage for use in therapeutic settings. Future work should aim to determine whether bacterial phylogeny can be used to predict bacterial virulence in clinical settings, as well as identifying high-throughput *in vitro* or *in vivo* methods which can allow for the more efficient testing of phage for use as therapeutics.

## Acknowledgments

The authors would like to thank Daniel Padfield and Pauline D. Scanlan for useful discussion and comments.

S.K.W. was supported by a studentship funded by the Biotechnology and Biological Sciences Research Council (BBSRC) South West Biosciences Doctoral Training Partnership (BB/M009122/1). R.M.I. and B.L. were supported by a Sir Henry Dale Fellowship jointly funded by the Wellcome Trust and the Royal Society (109356/Z/15/Z). A.B. is supported by NERC (NE/S000771/1) and NSF-NERC (DEB-1556444).

For the purpose of Open Access, the author has applied a CCBY public copyright licence to any Author Accepted Manuscript version arising from this submission. All data and scripts used for these analyses are available on GitHub (https://github.com/Sarah-K-Walsh/2026-In-Vivo-Phage-Efficacy).

## Author contributions

Conceptualisation: SKW, BL, AB

Methodology: SKW, RMI

Data curation: SKW

Investigation: SKW

Formal analysis: SKW, RMI

Supervision: BL, AB

Visualisation: SKW, RMI

Writing (original draft): SKW, RMI

Writing (review and editing): SW, RMI, BL, AB

