## Supplementary Text for "Bacterial phylogeny explains variation in virulence but not phage efficacy *in vivo*"

[+] Equal contribution

**Table S1: *Staphylococcaceae* samples metadata.** Clonal complex (CC) and sequence type (ST), where available, were determined using PubMLST. *S. aureus* strains missing an ST are coagulase-negative and therefore do not have an MLST scheme and the *S. aureus* strains with STs but no CCs are not similar enough to other strains to be assigned a CC. In 'source', 'JPP' is John-Paul Pirnay, 'EF' is Ed Feil, and 'GP' is Gavin Paterson.

| Isolate ID | Species | Host | Country | Year | CC | ST | NCBI BioSample ID | Source |
|---|---|---|---|---|---|---|---|---|
| 13S44S9 | *S. aureus* | *Homo sapiens* | BEL | 2012 | 8 | 8 | SAMN31484028 | JPP |
| 271Y | *S. kloosii* | *Eptesicus serotinus* | GBR | 2015 | --- | --- | SAMN31484029 | EF |
| 2111F7LW | *M. sciuri* | *Pteropus livingstonii* | GBR | 2017 | --- | --- | SAMN31484030 | EF |
| 27420LC | *S. simiae* | *Pteropus livingstonii* | GBR | 2016 | --- | --- | SAMN31484031 | EF |
| 2745SW | *S. nepalensis* | *Pteropus livingstonii* | GBR | 2016 | --- | --- | SAMN31484032 | EF |
| 82B | *S. caeli* | Environmental | ITA | 2013 | --- | --- | SAMEA2297795 | GP |
| 8325-4 | *S. aureus* | --- | --- | --- | 8 | 8 | SAMN31484033 | EF |
| AR03918O1 | *S. aureus* | *Sciurus carolinensis* | GBR | 2018 | --- | 133 | SAMN31484034 | EF |
| AR05S1 | *S. aureus* | *Sciurus carolinensis* | GBR | 2015 | --- | --- | SAMN31484035 | EF |
| AR05618O1 | *S. aureus* | *Sciurus carolinensis* | GBR | 2018 | --- | 49 | SAMN31484036 | EF |
| ASARM61 | *S. aureus* | *Homo sapiens* | GBR | 2006 | 22 | 22 | SAMN31484037 | EF |
| ASARM70 | *S. aureus* | *Homo sapiens* | GBR | 2006 | 22 | 22 | SAMN31484038 | EF |
| ASARM71 | *S. aureus* | *Homo sapiens* | GBR | 2006 | 22 | 22 | SAMN31484039 | EF |
| ASARM72 | *S. aureus* | *Homo sapiens* | GBR | 2006 | 22 | 22 | SAMN31484040 | EF |
| ASARM73 | *S. aureus* | *Homo sapiens* | GBR | 2006 | 22 | 22 | SAMN31484041 | EF |
| ASARM74 | *S. aureus* | *Homo sapiens* | GBR | 2006 | 22 | 22 | SAMN31484042 | EF |
| NCTC7692 | *S. saprophyticus subsp. saprophyticus* | Environmental | --- | 1948 | --- | --- | SAMEA3517999 | GP |
| NCTC11320 | *S. hominis spp hominis* | *Homo sapiens* | USA | 1975 | --- | --- | SAMEA3539708 | GP |
| NCTC11043 | *S. xylosus* | *Homo sapiens* | USA | 1975 | --- | --- | SAMEA3539705 | GP |
| B128S3 | *S. aureus* | *Sciurus carolinensis* | GBR | 2015 | 1 | 188 | SAMN31484043 | EF |
| B142S1 | *S. aureus* | *Sciurus carolinensis* | GBR | 2015 | --- | 692 | SAMN31484044 | EF |
| DAR04181C1 | *S. aureus* | *Cervus elaphus* | GBR | 2018 | 8 | 1958 | SAMN31484045 | EF |
| DAR06181LC1 | *S. aureus* | *Cervus elaphus* | GBR | 2018 | --- | 3237 | SAMN31484046 | EF |
| DAR091813 | *S. aureus* | *Cervus elaphus* | GBR | 2018 | --- | 425 | SAMN31484047 | EF |

| | | | | | | | | |
|---|---|---|---|---|---|---|---|---|
| DEU1 | *S. aureus* | *Homo sapiens* | TUR | 2009 | 8 | 239 | SAMN31484048 | EF |
| DEU2 | *S. aureus* | *Homo sapiens* | TUR | 2009 | 8 | 239 | SAMN31484049 | EF |
| DSM104441 | *S. edaphicus* | Environmental | ATA | 2013 | --- | --- | SAMN31484050 | GP |
| DSM107950 | *S. pseudoxylosus* | *Bos taurus* | FRA | 2002 | --- | --- | SAMN31484051 | GP |
| DSM18669 | *S. saprophyticus subsp. Bovis* | *Bos taurus* | CZE | 1996 | --- | --- | SAMN31484052 | GP |
| DSM21284 | *S. pseudo-intermedius* | *Felis catus* | BEL | 2008 | --- | --- | SAMN31484053 | GP |
| NCTC12218 | *S. schleiferi subsp. coagulans* | *Homo sapiens* | --- | 1988 | --- | --- | SAMEA3221103 | GP |
| DSM6628 | *S. schleiferi subsp. schleiferi* | *Canis lupus* | --- | 1991 | --- | --- | SAMN31484054 | GP |
| EOE23 | *S. aureus* | *Homo sapiens* | GBR | 1998 | 30 | 36 | SAMN31484055 | EF |
| EOE03 | *S. aureus* | *Homo sapiens* | GBR | 1998 | 30 | 3488 | SAMN31484056 | EF |
| EOE30 | *S. aureus* | *Homo sapiens* | GBR | 1998 | 30 | 36 | SAMN31484057 | EF |
| EOE35 | *S. aureus* | *Homo sapiens* | GBR | 2003 | 30 | 36 | SAMN31484058 | EF |
| EOE41 | *S. aureus* | *Homo sapiens* | GBR | 2005 | 30 | 36 | SAMN31484059 | EF |
| EOE42 | *S. aureus* | *Homo sapiens* | GBR | 2005 | 30 | 36 | SAMN31484060 | EF |
| HU25 | *S. aureus* | *Homo sapiens* | BRA | 1905 | 8 | 239 | SAMN31484061 | EF |
| JW32660O5 | *S. aureus* | *Sciurus carolinensis* | GBR | 2018 | --- | 49 | SAMN31484062 | EF |
| JW30866O BHY3 | *S. aureus* | *Sciurus carolinensis* | GBR | 2018 | --- | 49 | SAMN31484063 | EF |
| JW31330L BHY2 | *S. aureus* | *Sciurus carolinensis* | GBR | 2018 | --- | 49 | SAMN31484064 | EF |
| JW31330O BHY1 | *S. aureus* | *Sciurus carolinensis* | GBR | 2018 | --- | 49 | SAMN31484065 | EF |
| MU1 | *S. aureus* | *Homo sapiens* | TUR | 2010 | 8 | --- | SAMN31484066 | EF |
| MU2 | *S. aureus* | *Homo sapiens* | TUR | 2010 | 8 | 239 | SAMN31484067 | EF |
| NCTC11042 | *S. haemolyticus* | *Homo sapiens* | CZE | 1976 | --- | --- | SAMEA3233544 | GP |
| NCTC11046 | *S. simulans* | *Homo sapiens* | CZE | 1976 | --- | --- | SAMEA3504572 | GP |
| NCTC11047 | *S. epidermidis* | *Homo sapiens* | CZE | 1976 | --- | 5 | SAMEA3233545 | GP |
| P32 | *S. aureus* | *Homo sapiens* | POL | 1996 | 8 | 239 | SAMN31484068 | EF |
| SaTPS3026 | *S. aureus* | *Homo sapiens* | AUS | 2013 | 93 | 93 | SAMN31484069 | EF |
| SaTPS3043 | *S. aureus* | *Homo sapiens* | AUS | 2013 | 30 | 30 | SAMN31484070 | EF |
| SaTPS3072 | *S. aureus* | *Homo sapiens* | AUS | 2013 | 1 | 1 | SAMN31484071 | EF |
| SaTPS3097 | *S. aureus* | *Homo sapiens* | AUS | 2013 | 8 | 8 | SAMN31484072 | EF |
| SaTPS3104 | *S. aureus* | *Homo sapiens* | AUS | 2013 | 93 | 93 | SAMN31484073 | EF |
| SaTPS3105 | *S. aureus* | *Homo sapiens* | AUS | 2013 | 93 | 93 | SAMN31484074 | EF |
| SAR1018S1 | *S. aureus* | *Ovis aries* | GBR | 2018 | 8 | 8 | SAMN31484075 | EF |
| SAR1218N1 | *S. aureus* | *Ovis aries* | GBR | 2018 | --- | 1640 | SAMN31484076 | EF |

| | | | | | | | | |
|---|---|---|---|---|---|---|---|---|
| SAR1418N1 | *S. aureus* | *Ovis aries* | GBR | 2018 | --- | 130 | SAMN31484077 | EF |
| USFL008 | *S. aureus* | *Homo sapiens* | USA | 2009 | 8 | 8 | SAMN31484078 | EF |
| USFL009 | *S. aureus* | *Homo sapiens* | USA | 2009 | 8 | 8 | SAMN31484079 | EF |
| USFL012 | *S. aureus* | *Homo sapiens* | USA | 2009 | 8 | 8 | SAMN31484080 | EF |
| USFL016 | *S. aureus* | *Homo sapiens* | USA | 2009 | 8 | 8 | SAMN31484081 | EF |
| USFL018 | *S. aureus* | *Homo sapiens* | USA | 2009 | 8 | 8 | SAMN31484082 | EF |
| USFL020 | *S. aureus* | *Homo sapiens* | USA | 2009 | 8 | 8 | SAMN31484083 | EF |

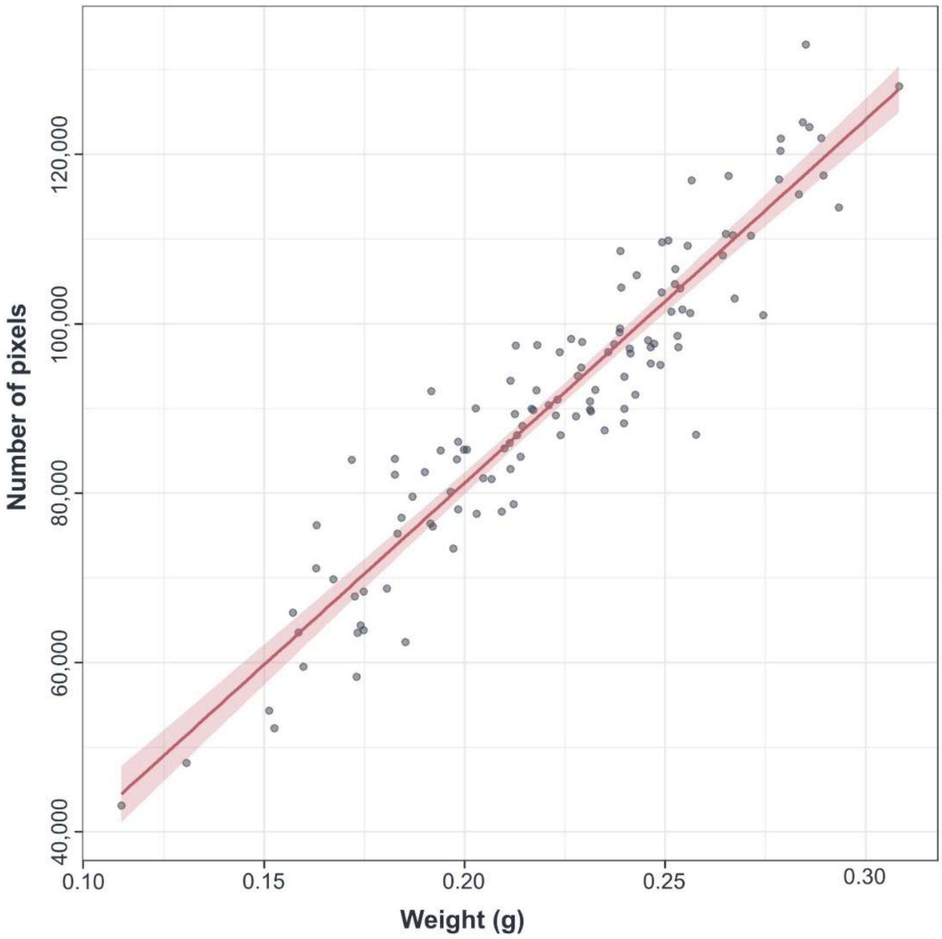


**Figure S1: Correlation between the weight of *G. mellonella* and the number of pixels they have when photographed**. 120 *G. mellonella* larvae were weighed and photographed to determine pixel area. The relationship between weight and pixel area was then assessed using a linear regression model and found to be strongly positively correlated (Pearson's $r = 0.94$, $p <0.001$).

A.

B.

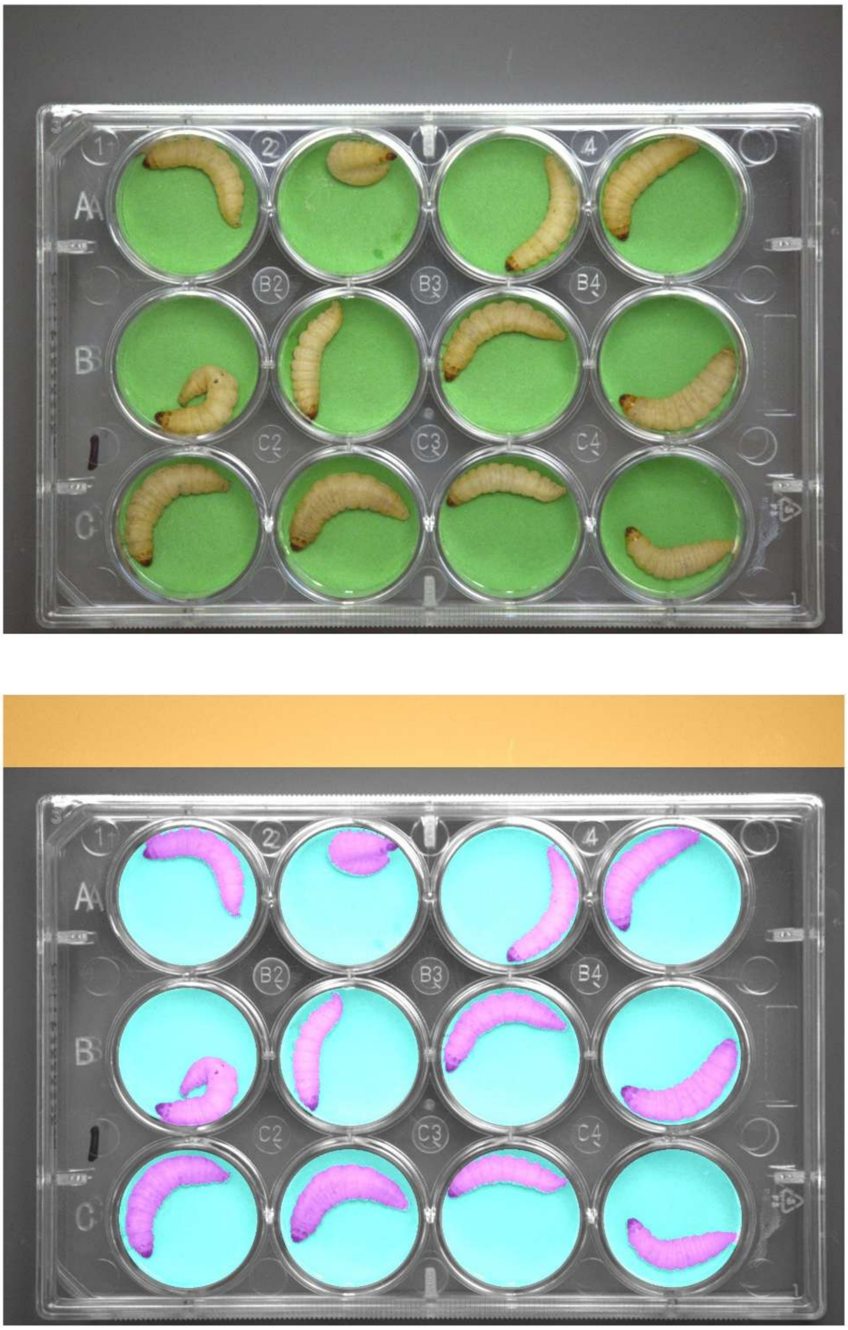

**Figure S2: Automated image segmentation algorithm.** Images show an example plate of inoculated *G. mellonella* larvae after the image has been demosaiced and white balanced (A), and then as a false colour validation image showing segmentation masks (B). In the false colour image, the orange area represents the rectangle of 18% neutral grey card used for white balancing, the blue areas represent the individual well masks, and the purple areas the individual larva masks.

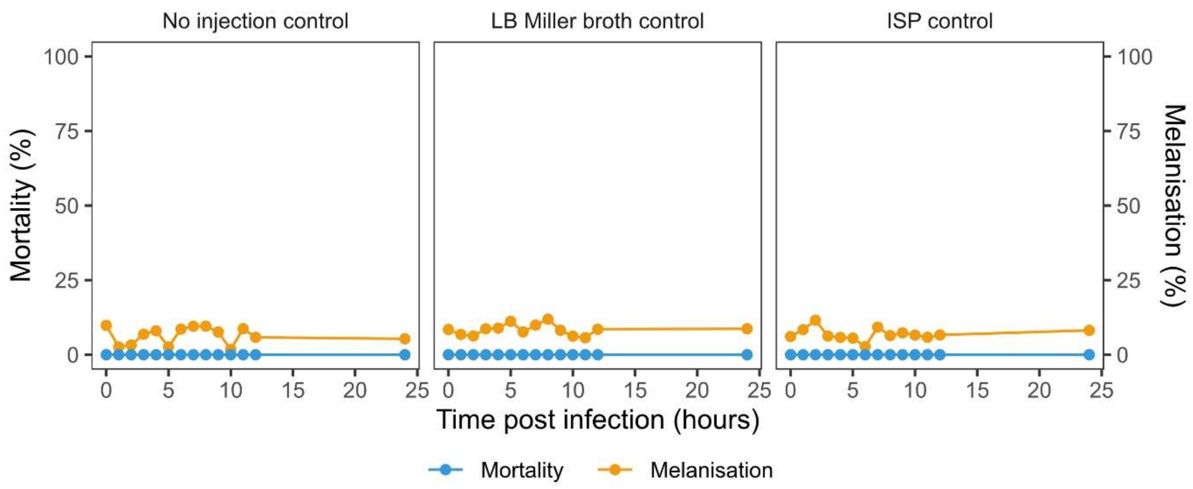


**Figure S3: Mortality and melanisation in *G. mellonella* infection controls.** Panels show the mean mortality and melanisation percentages of 120 *Galleria mellonella* larvae from each infection control treatment.

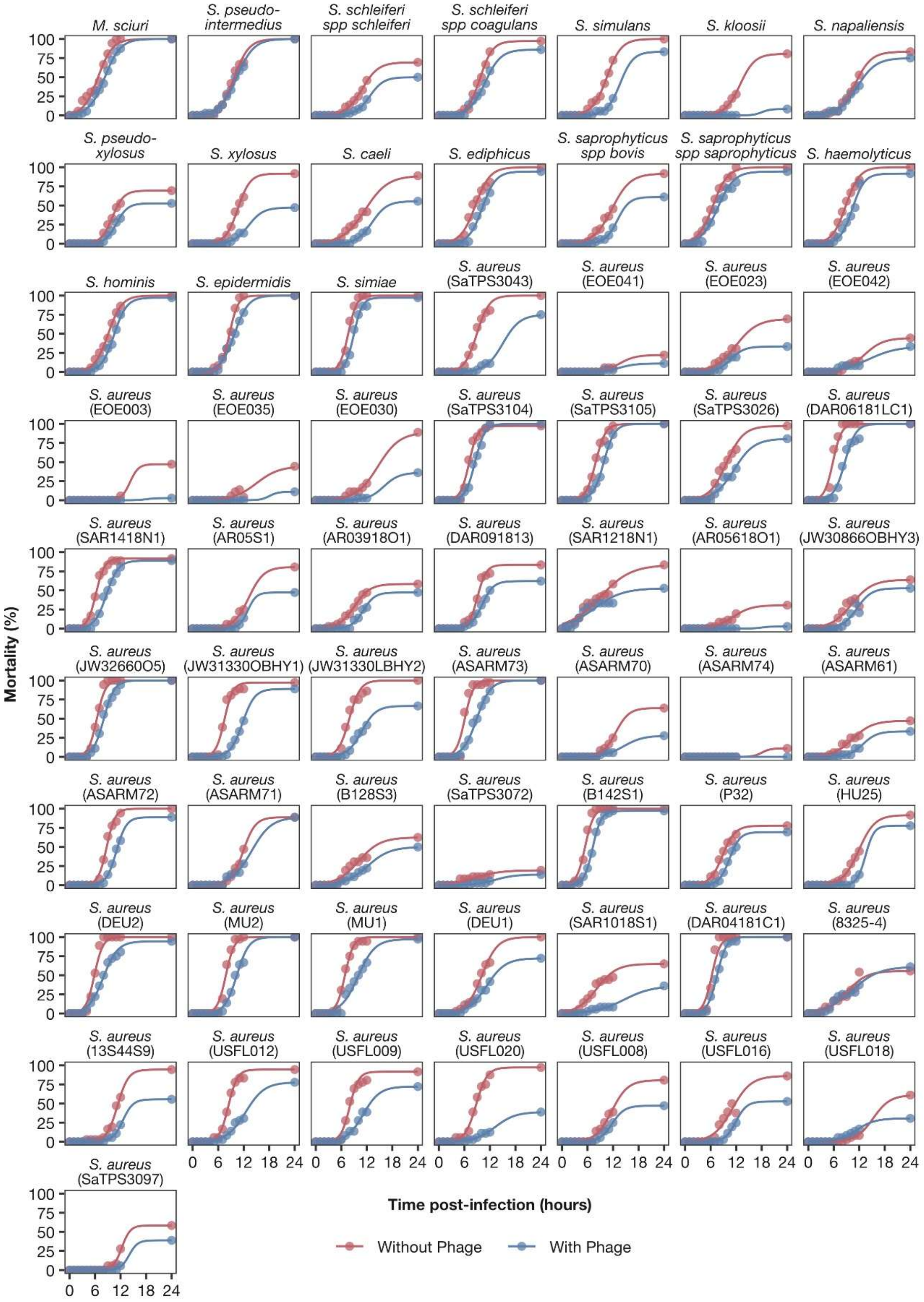


**Figure S4: Mortality curves of *G. mellonella* larvae infected with one of 64 *Staphylococcaceae* isolates in the presence or absence of bacteriophage, ISP.** Panels show the mean percentage mortality of *Galleria mellonella* larvae (solid lines) infected with an isolate of *Staphylococcaceae* both with (blue) and without (red) the bacteriophage ISP. Lines illustrate the non-linear least squares model fits for sigmoidal mortality curves under the “cured” assumption (i.e., where the endpoint observed mortality is considered the upper asymptote of each curve).

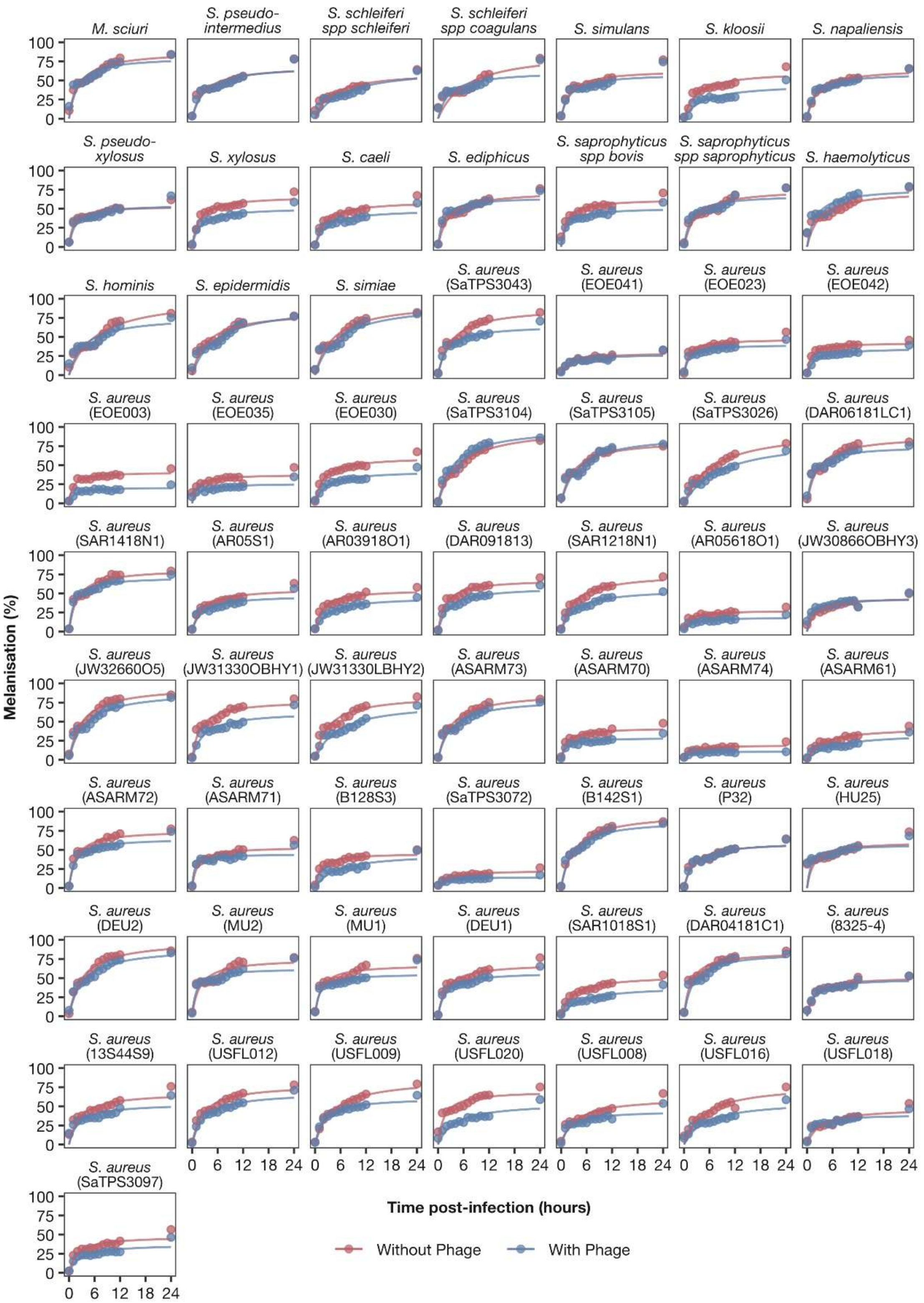


**Figure S5: Melanisation curves of *G. mellonella* larvae infected with one of 64 *Staphylococcaceae* isolates in the presence or absence of bacteriophage, ISP.** Panels show the mean percentage melanisation of *Galleria mellonella* larvae (solid lines) infected with an isolate of *Staphylococcaceae* both with (blue) and without (red) the bacteriophage ISP. Lines illustrate the non-linear least squares model fits for Michaelis-Menten curves.

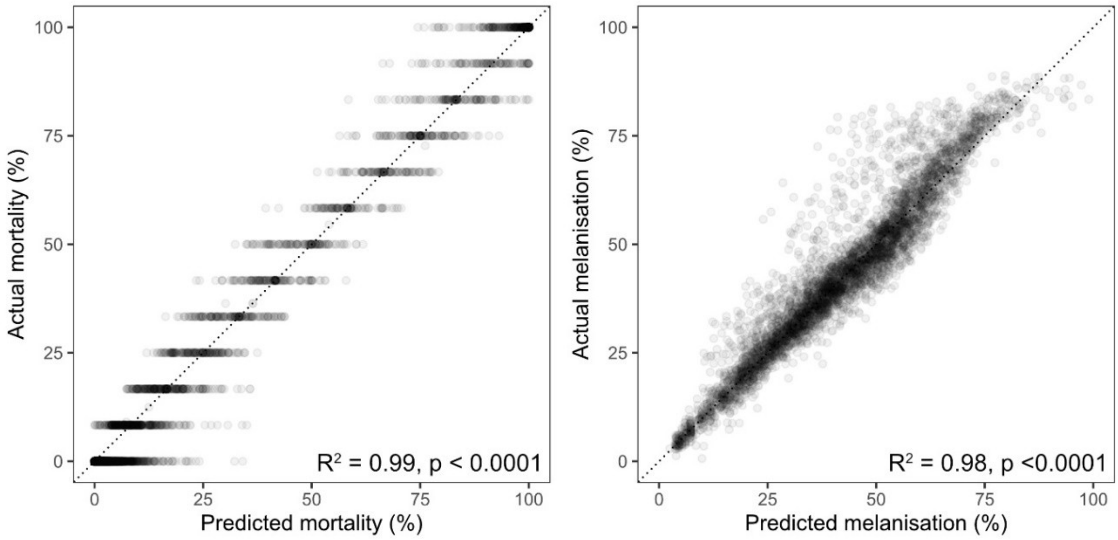


**Figure S6: Non-linear least squares model fits for mortality and melanisation data.** Plots show the relationship between model predicted (x axis) and observed (y axis) values for mortality and melanisation. Predicted values are taken from non-linear least squares models of sigmoidal curves under the "cured" assumption (mortality, left-hand plot) or Michaelis-Menten curves (right-hand plot). $R^2$ values and significances of the correlation between predicted and observed values are taken from linear regression models.

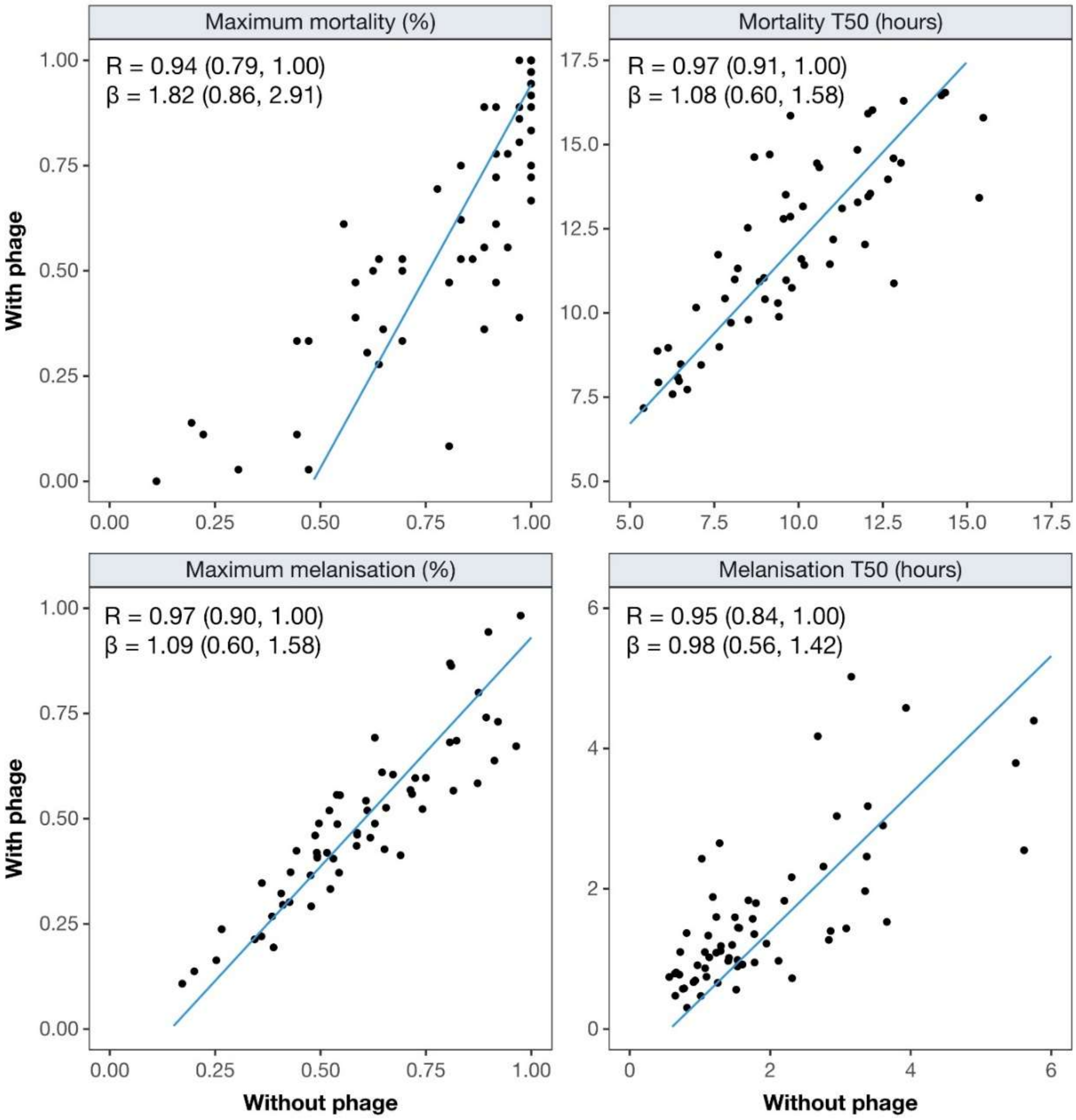


**Figure S7: Inter-strain correlations between mortality and melanisation parameters in the presence and absence of ISP.** Plots show the inter-strain correlations of the 'With phage' and 'Without phage' treatments for each mortality and melanisation parameter, with points indicating the mean value of each *Staphylococcaceae* strain. Trendlines, correlation coefficients and slopes are taken from phylogenetic mixed models of structure (1), with trendlines whose coefficient and slope estimates do not include zero highlighted in blue.
